# Site-specific phosphorylation of histone H1.4 is associated with transcription activation

**DOI:** 10.1101/814129

**Authors:** Ankita Saha, Christopher Seward, Lisa Stubbs, Craig Andrew Mizzen

## Abstract

Core histone variants like H2A.X and H3.3 and their modified forms serve specialized roles in chromatin processes that depend on their genomic distributions and their interaction with chromatin components. Similarly, previous evidence from our lab and others suggest that amino acid sequence variant forms of the linker histone family and specific posttranslational modifications on these variants also result in distinct functions. These inferences are contrary to the notion that the H1 family function as redundant repressors. Here, we provide the first genome-wide evidence that when phosphorylated at a specific C-terminal domain site i.e serine 187, the linker histone H1.4 is enriched at active promoters. This is in direct contrast to previous reports that suggest that phosphorylation of H1 leads to their dissociation from chromatin. Using a highly specific pS187H1.4 antibody earlier developed in the lab, we studied the distribution patterns of pS187H1.4 in estradiol-responsive MCF7 cells where we demonstrated the inducible nature of this modification. We also used public MCF7 data to confirm the association of pS187H1.4 with well-known active transcription marks. These data suggest that linker histones and their modified forms have a more nuanced role than previously understood and may even play a role in transcription regulation.

## INTRODUCTION

The histone H1 family, also known as the linker histone, is a key structural component of nucleosome and higher order chromatin organization. They are present in non-allelic amino acid sequence-variant forms in many metazoans[1]. Humans express eleven H1 variants; seven variants are ‘somatic’; whereas the rest are selectively expressed in germline tissues. Five of the seven somatic variants-H1.1, H1.2, H1.3 H1.4 and H1.5 - are replication-dependent and are expressed predominantly during the S-phase of the cell cycle. These variants contain a more conserved amino acid sequence as compared to the replication-independent variants, namely H1.0 and H1.X[2]. Metazoan H1 variants share a common tripartite structure containing a central globular domain (GD) flanked by a short N-terminal domain (NTD) and a long C-terminal domain (CTD). Fluorescence recovery after photo-bleaching (FRAP) analyses have shown H1 binding to chromatin is dynamic in nature [3, 4] and the differences in the CTDs of these variants contribute to their varying binding affinities [5]

H1 CTDs and NTDs undergo several types of posttranslational modifications amongst which phosphorylation was observed to be the most abundant modification. Studies showed that specific sites within the CTD were phosphorylated during interphase and mitosis. During interphase, the serine-containing consensus CDK motifs (SPXZ-S: Serine P: Proline X: amino acid Z: Lysine (L)/arginine (R)) were the predominant sites of phosphorylation. These sites are also phosphorylated during mitosis, but during that time, threonine-containing CDK sites (TPXZ) and some non-CDK motifs are also phosphorylated [6, 7].

Early studies from our lab, conducted in HeLa S3 cells, identified the predominant interphase phosphorylation sites in H1.2 and H1.4 as H1.2-S173, H1.4-S172 and H1.4-S187 [8]. Using novel, highly specific antisera developed for H1.4 phosphorylated at S187 (pS187-H1.4), we showed that pS187-H1.4 is enriched at sites of transcription by RNAP I and II. In a more recent study, we showed that the global levels of H1 phosphorylation at H1.5-Ser18 (pS18-H1.5), H1.2/H1.5-Ser173 (pS173-H1.2/5) and pS187-H1.4 are subject to differential regulation and that CDK9 phosphorylated pS187-H1.4 was associated with maintenance of pluripotency [9]. This provided the first evidence that pS187-H1.4 is may be important for transcriptional activation.

Here, we provide the first genome-wide view of pS187-H1.4 dynamics using an inducible system – specifically, estradiol-responsive MCF7 cells. Using an affinity purified version of the previously generated pS187-H1.4 antibody, we show that pS187-H1.4 is strongly associated with the promoters of ‘active’ genes. Further, we show that these ‘active’ signals are quenched by a specific CDK9 inhibitor, Flavopiridol, resulting in gene repression. We combined pS187-H1.4 ChIP-sequencing data in conjunction with publicly available data for RNA Polymerase II (RNAP II) conducted in a similar system to show that pS187-H1.4 co-localizes with active RNAP II peaks. Taken together, these data provide evidence towards a more nuanced role of H1 in and puts forth the possibility of a new layer of regulation within the accepted model of transcriptional activation.

## MATERIAL AND METHODS

### Cell culture

MCF7 cells, a kind gift from Dr. Katzenellenbogen (UIUC), were grown in RPMI 1640 media supplemented with 5% fetal bovine serum (FBS) and 1% Penicillin-streptomycin and subcultured by 0.25% trypsin-EDTA treatment followed by seeding at 1.5× 10^6^ cells per 10cm plate. Before use, the cells were subjected to 72 hours estradiol depletion by growing in phenol-red free RPMI1640 supplemented with charcoal dextran treated FBS (CD-FBS). Following hormone depletion, the cells were induced with 20nM beta-estradiol (E2) dissolved in ethanol for 30 minutes before harvesting. In case of drug treatments, the cells were pre-treated with Flavopiridol (NIH AIDS Reagent Program) for 1Hr before E2 induction to selectively inhibit CDK9.

### siRNA knockdown

siRNA treatments were conducted using Lipofectamine RNAiMAX (Invitrogen) and H1.2/H1.4, H1.4, CDK9, CDK7 siRNA according to manufacturer’s protocol. The cells were seeded the day prior to transfection to achieve 50-60% confluency at the time of treatment. siRNA and lipofectamine were diluted in Opti-MEM (Gibco), mixed, incubated for 5 mins and the complexes were then added directly to the cells now growing in phenol-red free, CD-FBS Media to allow estradiol depletion along with siRNA knockdown simultaneously. After 72 hours, the cells were E2/ ethanol for 30 minutes before harvesting.

### Chromatin Immunoprecipitaiton

Chromatin immunoprecipitation was performed using protocols described before [9] with minor adjustments. Briefly, cells were cross-linked using final concentration 1% methanol-free paraformaldehyde (Thermo Fisher Scientific) for 8 minutes quenched by 125mM final concentration glycine for 10 minutes. The cells were then washed 3 times with cold PBS before scraping and resuspension in ChIP lysis buffer supplemented with protease and phosphatase inhibitors. Chromatin was then sonicated using 30sec on/off cycles for 25 minutes in the BiorupterTM UCD-200 (Diagenode, Liège, Belgium) sonicator to a mean sheared length of ~500kb. Following centrifugation, the supernatants were diluted tenfold using ChIP-dilution buffer. 1ml aliquots representative of 1.5-2× 10^6^ cells were then incubated with primary antibody overnight at 4°C. The antibody-chromatin complexes were then incubated with 50μl BSA-blocked Dynabeads (Invitrogen) for 4H at 4°C and then collected using a magnetic separator. Following sequential ChIP wash-buffer and TE buffer washes, the beads were eluted using 200μL freshly prepared elution buffer. 200mM final concentration NaCl was added to these eluates and incubated at 65°C overnight to reverse crosslink the immunocomplexes. Following RNAse A and Proteinase K digestion, the DNA was then purified using phenol/chloroform extraction and then precipitated using glycogen as a carrier. The precipitated DNA was then dissolved in Tris-EDTA buffer and used for qRT-PCR or Illumina sequencing.

### ChIP-Western blot

MCF7 cells (+/−E2) were collected following trypsinization. The nuclei were isolated for chromatin extraction. Briefly, the cells were washed with 0.1% tween-20 in 1XPBS, 0.1%NP40 in Tris-MgCl2 (TM2) buffer, and TM2 buffer to extract the nuclei. The nuclei were then subjected to micrococcal nuclease (2 units) treatment for 6 minutes before using 0.5X PBS for overnight chromatin extraction. pS187-H1.4 antibody was then added to this chromatin extract and allowed to bind overnight. Sepharose beads were used to elute the chromatin-antibody complex. This mixture was then boiled with Lammeli buffer and then run on a 4-20% polyacrylamide gel for western blot.

### Quantitative Real Time-PCR (qRT-PCR)

For expression analyses, RNA was extracted from cells using the trizol extraction method[10] and further purified using the Qiagen DNA purification kit. cDNA was prepared using the Superscript III first strand synthesis system (Thermo Fisher Scientific) according to manufacturer protocol.

cDNA or ChIP products were used for qRT-PCR with SYBR-Green Mix (Applied Biosystems) and primers listed in supplementary table 1.

### Sequencing and Bioinformatic analysis

ChIP products were used to prepare libraries using a kit from KAPA Biosystems. Manufacturer protocols were followed for sheared DNA size 500-700Kb. Quality control of libraries was conducted using a bioanalyzer and then sequenced using the Illumina-Hiseq 4000 by the W. M. Keck Center for Comparative and Functional Genomics at the Roy J. Carver Biotechnology Center (University of Illinois). Sequence data were mapped with Bowtie2 [11] to the UCSC hg19 genome, using default settings. Mapped sequence data were analyzed for peaks using HOMER (Hypergeometric Optimization of Motif EnRichment) v4.10 [12]. Samples were converted into tag directories, and QC was performed using read mapping and GC bias statistics. Histone peaks were then called from the Tag Directories with default factor settings, except local filtering was disabled (-L 0), peak size was set at 200bp (-size 200) and minimum distance between peaks was set at 150 (-min dist 150), to increase the sensitivity of the peak calling and identify individual subunits of multi-histone peaks. After peak calling, peak files were annotated to the human hg19 genome using HOMER’s annotation script to assign peaks to nearest genes, and associate peaks with estrogen responsive differential genes identified elsewhere by Gro-Seq[13]. BigWiggle pileup files were generated using HOMER’s makeBigWig.pl script with default settings (normalized to 10m reads) and uploaded to a UCSC Genome Browser track hub for visualization. Differential chromatin peaks were identified using the HOMER getDifferentialPeak.pl script, looking for any peaks that changed at least two-fold between conditions with a significance cutoff of 1 × 10-4. Genes annotated 100kb from differential H3K27ac peaks were submitted for GO analysis to DAVID and GREAT [14, 15]. Metagenome profiles were generated using the HOMER makeMetaGenomeProfile.pl script using default settings. Aggregate plots were generated using the HOMER annotatePeaks.pl script with size set to 4000-6000bp and binning set to 10bp.

Two biological replicates of pS187-H1.4 and pan-H1.4 ChIP-sequencing were performed. The IP efficiency of replicate 1 was better than replicate 2, however, the peak enrichment patterns agreed closely and correlation coefficient scores calculated between the replicates were >0.94. Browser views demonstrated close to identical trends (Supplementary Figure. 8). Data from the first replicate was used to generate plots for better visualization.

## RESULTS

### Genome wide distribution of pS187-H1.4 displays distinct patterns of enrichment

We previously generated a collection of unique, highly specific antisera using synthetic phospho-peptides and recombinant proteins and demonstrated their ability to recognize phosphorylation at single sites [8, 9]. These sites are either exclusive to an individual human H1 variant or are shared between only two variants. In addition, we also raised ‘pan H1’ antisera using whole recombinant H1 variant protein that does not distinguish between phosphorylated forms [8]. While the phospho-H1 antisera provide a measure of phosphorylation between specific sites, the pan-H1 antisera provides a comparison of the H1 variant amounts present, regardless of its phosphorylation status. Evidence of the specificity of pS187-H1.4, pan-H1.4, pS18-H1.5 and pan-H1.5 has been described previously [8, 16].

Although these antisera are highly specific, their efficiency for high throughput studies was limited. Therefore, for this study, we purified pS187-H1.4 and pan-H1.4 antisera using a modified affinity chromatography approach to generate a highly efficient set of H1 antibodies. The purified pS187 H1.4 antibody was re-tested for specificity in accordance to ENCODE guidelines as shown in supplementary figures 1-3. We chose MCF7 cells because they have been extensively studied, and therefore a large amount of high-throughput data are available on public platforms for comparison to and further analysis of our data [13, 17, 18].

We used the purified pS187-H1.4 and pan-H1.4 antibodies to perform chromatin immunoprecipitation (ChIP) on MCF7 cells to reveal the genome-wide distribution of all versions of H1.4 protein (pan-H1.4) and the specifically phosphorylated version (pS187-H1.4). Peaks generated from the sequencing data were then associated with their nearest genes. We immediately noted a special enrichment of pS187-H1.4 ChIP peaks around promoter regions (Fig. 1A). The association of pS187-H1.4 peaks with promoters genome wide was significant, with a log ratio of 2.85 (Table1) when analyzed by the HOMER software (http://homer.ucsd.edu) [12]. In striking contrast, we observed that the pan-H1.4 signal was depleted at promoters. This pattern of pan-H1.4 localization has been previously reported using another antibody [19], providing independent corroboration of our data.

**Figure 1.**
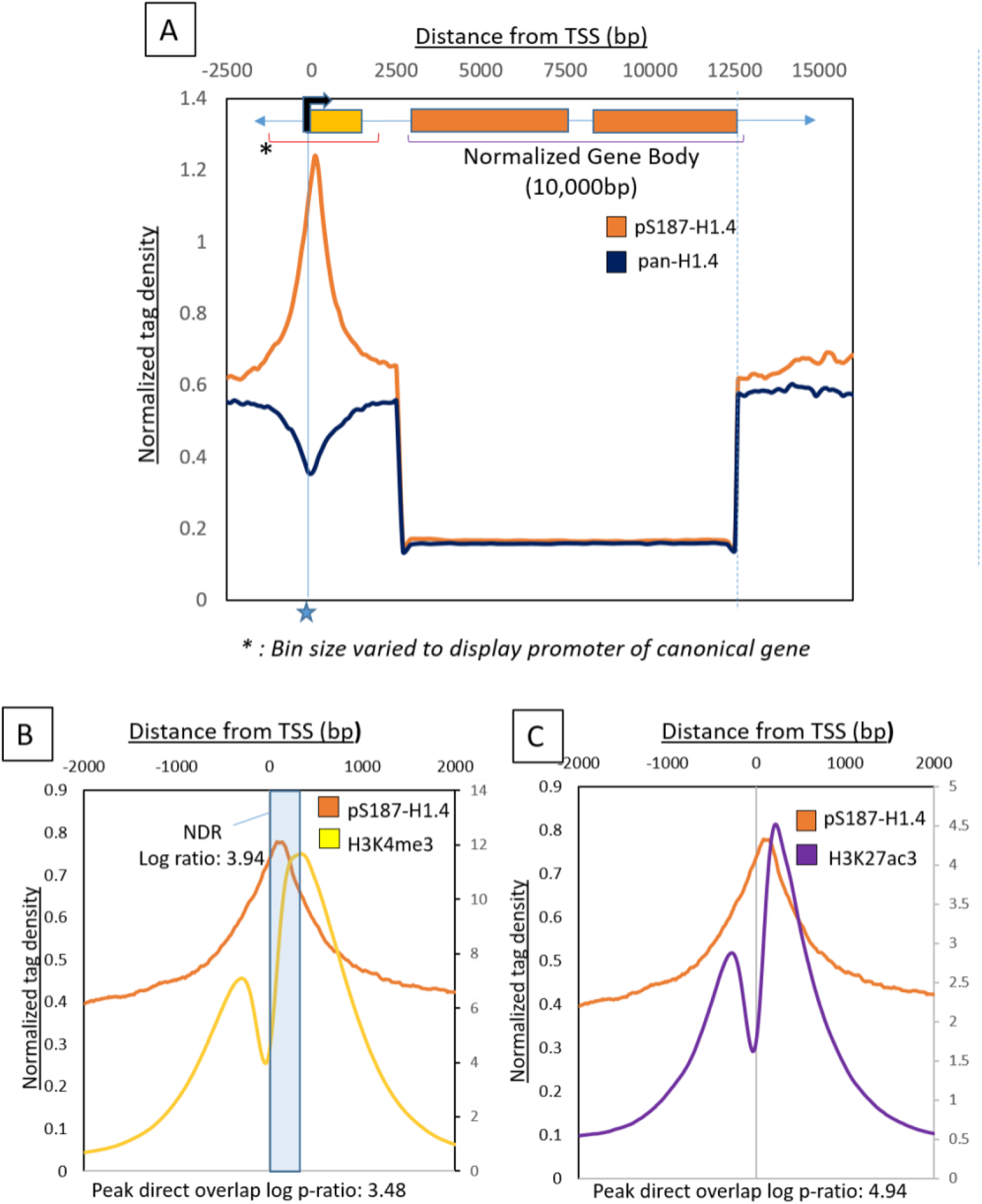
Global distribution of pS187-H1.4 and pan-H1.4 in MCF7 cells Figure 1 (A): A metagenome profile generated with pS187-H1.4 and pan-H1.4 ChIP-sequencing data to study enrichment across a typical gene. The gene body was mathematically defined and normalized to 10kb. +2.5kb and −2.5kb regions relative to the promoter were binned at 100bp. 2.5kb to 10kb represents a normalized gene body binned at 200bp. Typical transcription start sites (TSSs) marked by a star showed maximum enrichment of the pS187-H1.4 signal (orange trace). There was a depletion of the pS187-H1.4 and the pan-H1.4 (navy blue trace) signals along the gene body. (B): Aggregate plot centered on the promoter showing average pS187-H1.4 signals and its overlap with H3K4me3 signals. The signal was aligned +2Kb and −2Kb relative to the Refseq TSSs H3K4me3 was plotted on a secondary axis (label on right side). The direct overlap of peak was calculated as a log ratio of 3.48. H3K4me3 flanked predicted NDR region is highlighted in blue. The overlap of the pS187-H1.4 with this NDR region is 3.94. (C): Aggregate plot centered on the promoter showing average pS187-H1.4 signals and its overlap with H3K27ac signals. H3K27ac was plotted on a secondary axis (label on right side). The direct overlap of peak was calculated as a log ratio of 4.94.

**Table 1:**
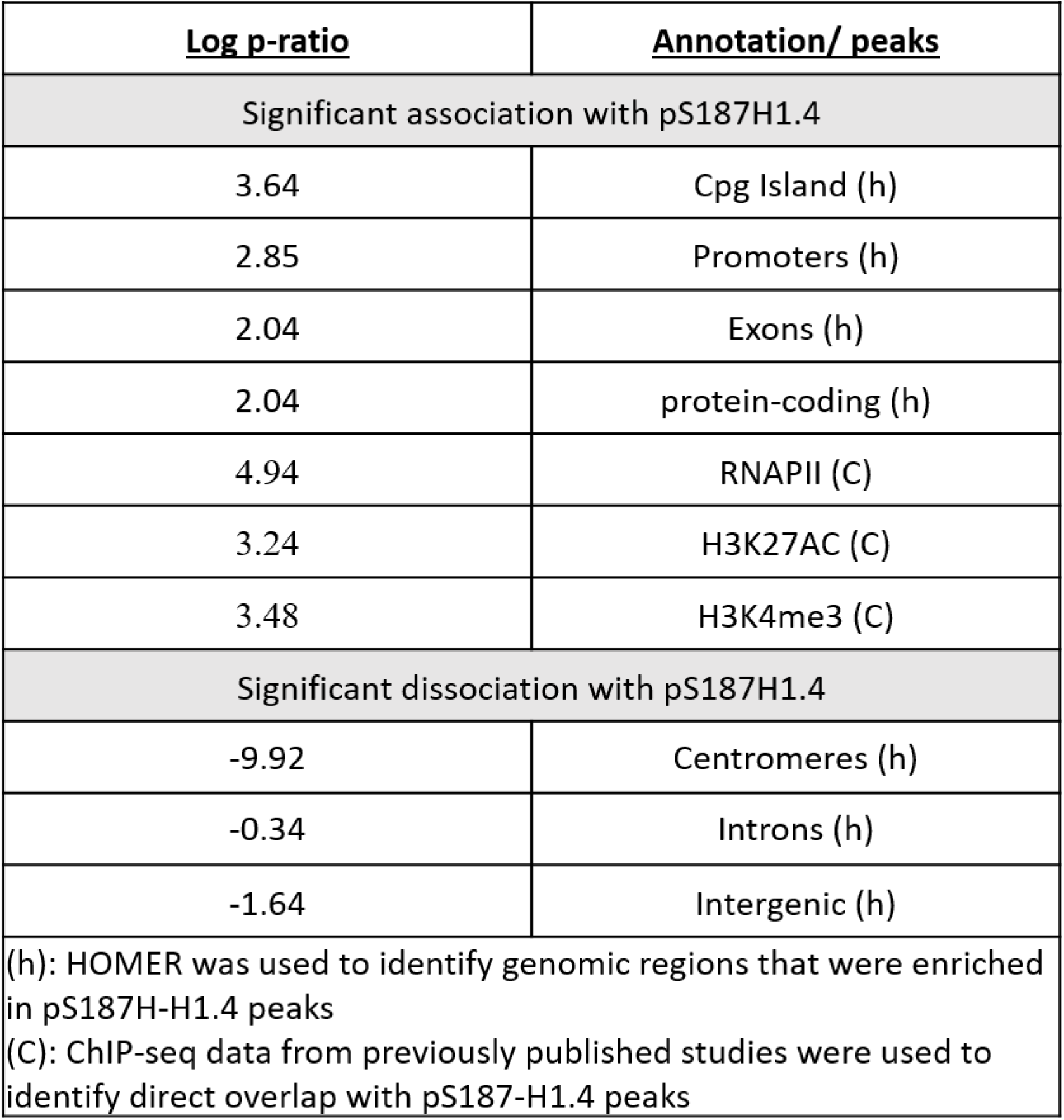
Summary table of pS187-H1.4 peak correlations with genomic regions and ‘active’ genomic marks.

In order to further characterize the status of the promoters associated with pS187-H1.4 peaks, we used H3K4me3 ChIP-Seq data from a separate study conducted on MCF7 cells [20]. The H3K4me3 epigenetic mark has widely been associated with ‘active’ transcription state or the ‘transcription readiness’ of nearby promoters [21, 22]. We plotted the average pS187-H1.4 and H3K4me3 signal around all promoters into an aggregate plot to study the trend of enrichment (Figure 1B). As illustrated by this figure, the pS187-H1.4 peaks overlap significantly with H3K4me3 peaks (Log p-ratio: 3.48). This suggests that pS187H1.4 peaks associate with ‘active’ or ‘poised’ promoters[23]

This alignment also revealed that pS187-H1.4 peaks align with the ‘dip’ between paired H3K4me3 peaks [24] identified as a H3 depleted or a ‘Nucleosome Depleted Region’ (NDR) [25]. A statistical analysis showed that the overlap between pS187H1.4 peaks and the H3K4me3-associated NDR was highly significant (log p-ratio: 3.48). In fact, it is 1.82 times more likely that a pS187-H1.4 peak will be found within this NDR than to overlap with the H3K4me3 reads *per se*. Confirming that the pS187H1.4 peaks were associated with ‘active’ promoters, we also found significant overlap with another ‘active’ promoter mark; H3K27ac [20] (Figure 1C) (log ratio: 4.94) thereby providing multiple points of corroboration.

### pS187-H1.4 peaks co-localize with promoter associated RNAPII

Based on the promoter-enriched location of the pS187-H1.4 peaks, we then analyzed the overlap of RNA Polymerase II (RNAPII), which is usually found to be proximal to the promoter of genes that are transcriptionally ready or actively transcribed. We used MCF7 ChIP-seq data from a different study [13] to generate aggregate plots for the RNAPII signal, and compared that signal with the average signal of pS187-H1.4 (Figure. 2A). This comparison revealed a significant direct overlap between the locations of pS187-H1.4 and RNAPII proteins at promoters in MCF7 cells (log ratio: 4.9). We also analyzed the overlap between RNAPII and pS187-H1.4 peaks throughout the genome and confirmed a positive correlation between the two (correlation R^2^ value 0.55; Figure 2B).

**Figure 2:**
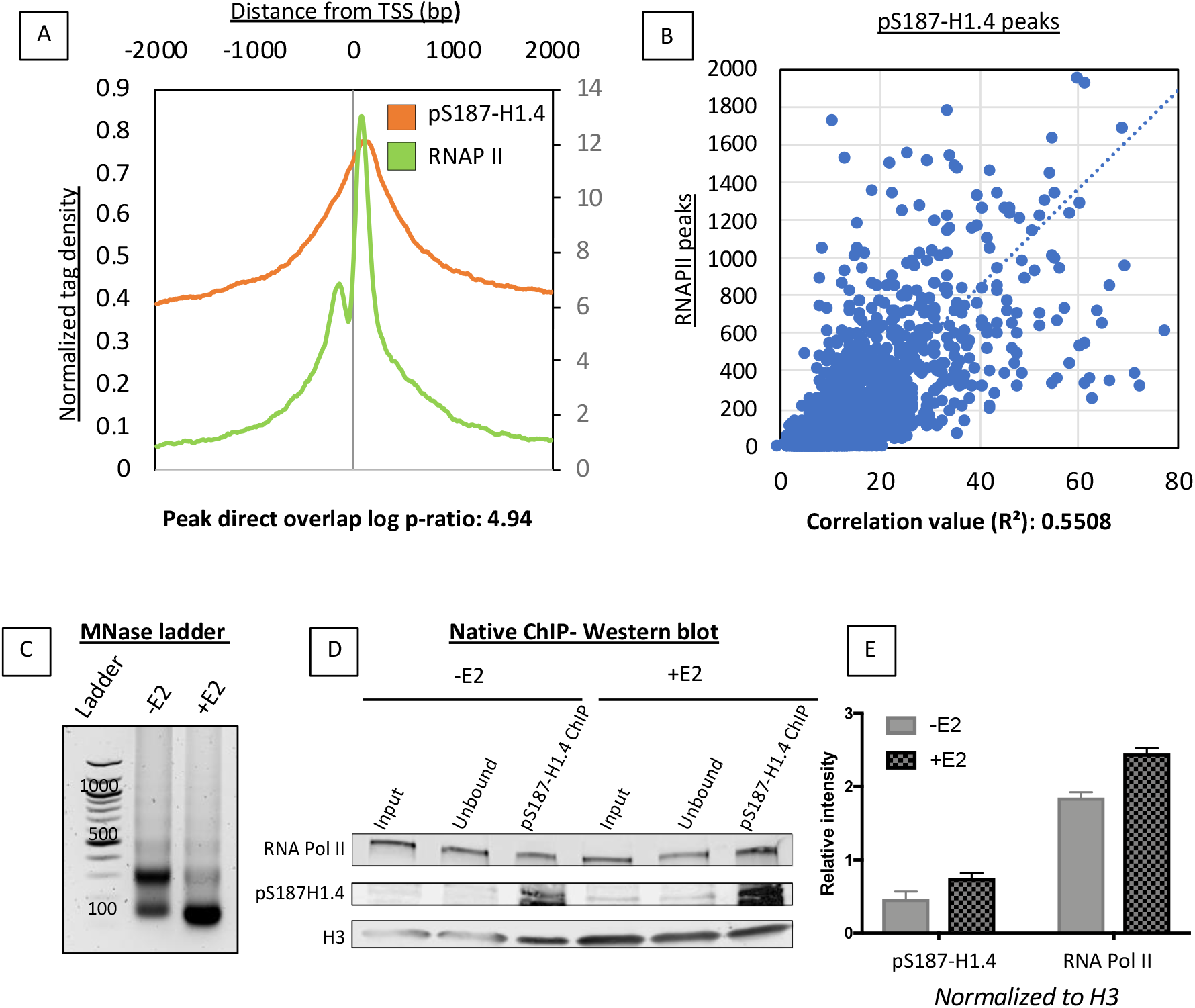
pS187-H1.4 associates with RNAPII. (A) Aggregate plot comparing average RNAPII (Green) and p187-H1.4 (Orange) peaks near promoter regions. The signal was aligned +2Kb and −2Kb relative to the Refseq TSSs. RNAPII plotted on a secondary axis (label on right side of graph). Direct overlap ratio calculated as log ratio: 4.94. (B) Correlation scatter plot of pS187-H1.4 and RNAPII peak overlap throughout the genome. The correlation co-efficient R_2_ calculated at 0.558. (C) 6 minute MNase digested DNA from estradiol untreated (-E2) and estradiol treated (+E2) MCF7 nuclei run on a 1.8% agarose gel. (D) Western blot following pS187-H1.4 native-ChIP. RNA pol II, pS187-H1.4 antibodies used for detection and H3 used as a loading control. (E) Relative intensity quantification of western blot in fig. 2D. H3 signal was used to normalize pS187-H1.4 and RNAP II signals.

In order to confirm the physical interaction of the pS187H1.4 with RNA pol II in our estradiol inducible MCF7 system, we performed native-ChIP, which does not rely on formaldehyde cross-linking, thereby providing an accurate and unbiased view of protein-protein interactions. Further, to ensure that the interaction between the RNAPII and the pS187H1.4 is within very close range ~1-2 nucleosomes, we used micrococcal nuclease digestion to obtain a majority of mononuclesomes (Figure.2C). Following chromatin immunoprecipitation with our pS187H1.4 antibody, we analyzed the samples using a western blot. Here we see that pS187H1.4 is enriched following estradiol stimulation. Strikingly, we also see that the pS187H1.4 pulls down RNAPII thereby demonstrating a stable physical interaction between the two most likely within the same nucleosome. This pulldown is enriched as a result of estradiol stimulation (Figures. 2D and 2E).

Taken together, these data show that pS187-H1.4 peaks have a distinct pattern of distribution and strong association with ‘active’ promoters. In order to better understand the genome-wide dynamics of pS187-H1.4 at the promoters, we sought to further explore the pS187-H1.4 binding in a transcription activating-estradiol- inducible system.

### Estradiol (E2) stimulation resulted in pS187-H1.4 peak enrichment

To ask whether pS187-H1.4 binds to promoters as they are activated, we used the purified pS187-H1.4 and pan-H1.4 antibodies to perform chromatin immunoprecipitation (ChIP) on MCF7 cells treated with 20nM 17β-Estradiol (E2) for 30 minutes. This treatment is sufficient to ‘activate’ the early estrogen-regulated network of genes, following 72 hours of hormone starvation [26].

The estradiol response can vary in different cell lines to the point where some genes may be oppositely regulated [27]. For example, genes involved in cell proliferation - like CDC2, CDC6, and Thymidine kinase 1 - are induced by E2 in MCF7 cells, whereas in the similar E2 responsive breast cancer cell line, 231ER+, these same genes are repressed [28]. The differences are likely due to the presence of different transcription factors and/or co-regulators in each cell type. To avoid such gene expression discrepancies we used data from independent studies but limited only to MCF7 cells [29].

We conducted ChIP-seq in both untreated and E2-treated MCF7 cells with pS187-H1.4 and pan-H1.4 antisera, and processed the data as described in the methods section. We then used the peaks from pan-H1.4 and pS187-H1.4 before and after E2 treatment to generate metagenome profiles to visualize their distribution across the genome with respect to a normalized gene (Supplementary figure. 6A). We aligned the 2013 pS187-H1.4 peaks from untreated cells and compared their distribution with 2830 peaks after E2 induction. These data showed that E2-induced pS187-H1.4 peaks show increased levels of promoter enrichment compared to the pS187-H1.4 peaks in untreated controls. The pattern of pS187-H1.4 distribution was otherwise similar before and after hormonal induction. Furthermore, the pattern of pan-H1.4 depletion at promoters and across gene bodies remained constant before and after E2 treatment. There was only a small overall increase in pS187H1.4 signal after E2 induction (as represented in a metagenome profile) when visualized in the context of all occupied genes, as expected since only ~15% of all genes are estradiol-responsive [13, 17, 30, 31]. Nevertheless, an increase in overall pS187-H1.4 promoter occupation was observed.

We focused our attention on promoter regions specifically in further analyses. First, we plotted the average pS187-H1.4 signal before and after E2 treatment, focused on 3000bp regions upstream and downstream of genome-wide TSSs (Figure 3A). This plot illustrates clearly that pS187H1.4 is most highly enriched at a position located approximately 150bp from transcription start sites, i.e. within the +1 nucleosome, where previous reports have found ‘paused’ polymerases to be bound [32, 33]. We also generated aggregate plots of pS187-H1.4 peaks in E2-treated cells to test overlap with H3K4me3, RNAPII and H3K27ac and enrichment of pS187-H1.4 peaks at ‘active’ promoters (Supplementary Figure 6 B, C&D). The overlap ratios were significant with log ratios of 3.25, 4.99 and 3.14 respectively with each ‘active’ mark. Taken together, these data show for the first time that pS187-H1.4 is associated with ‘active’ promoters and is suggestive of an interaction with transcriptional machinery.

**Figure 3:**
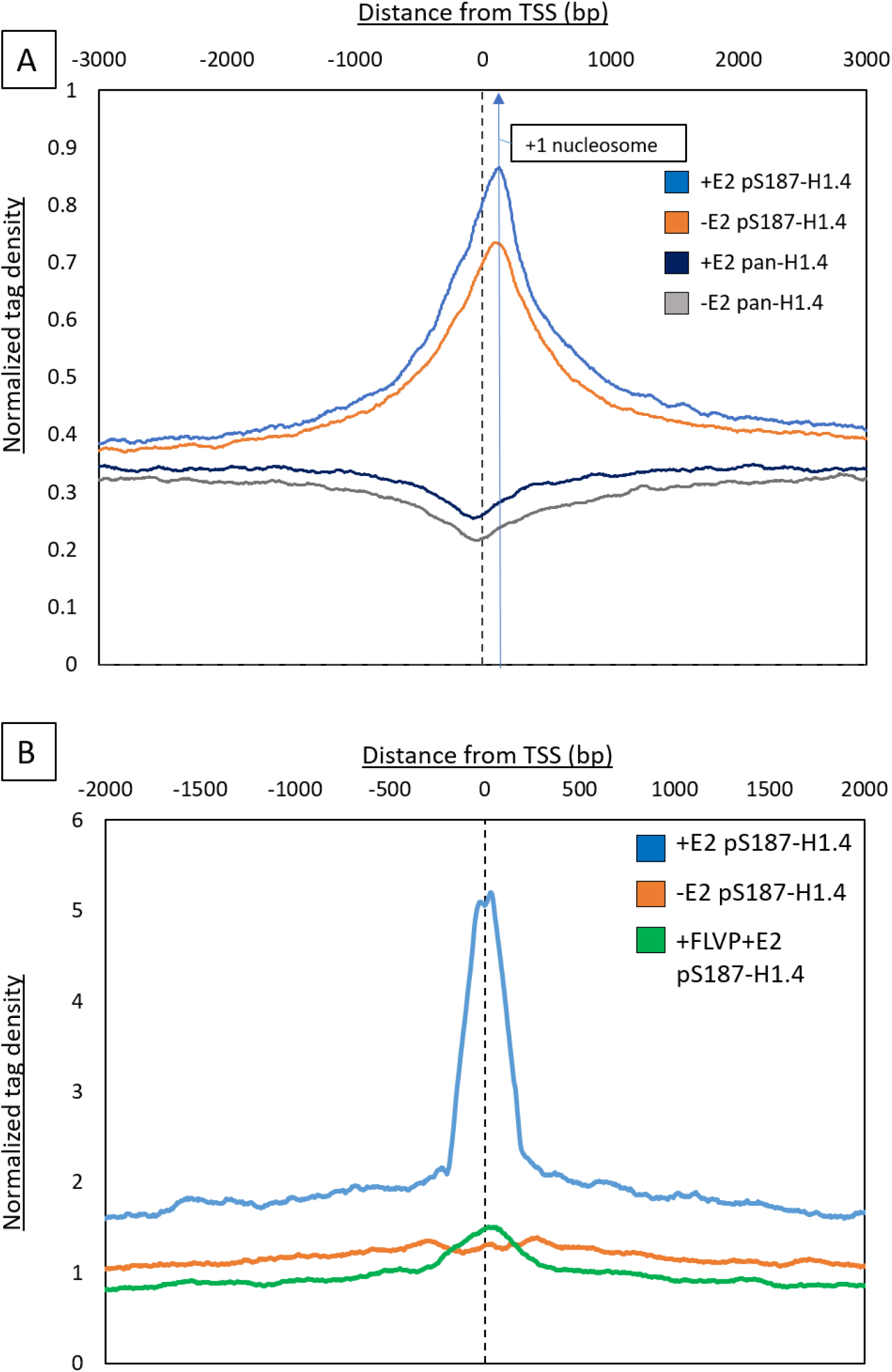
Estradiol induced pS187-H1.4 peaks near promoter. (A) Aggregate plot with average pS187H1.4 peaks before (Orange) and after (Blue) estradiol (E2) treatment. The signal was aligned +3Kb and −3Kb relative to the Refseq TSSs Following estradiol treatment, pS187-H1.4 is enriched. Highest signal of pS187-H1.4 shifted towards the predicted location of +1nucleosome. Pan-H1.4 before (Grey) and after (Navy blue) estradiol treatment is also seen. Depletion of signal at promoters is seen. (B) Aggregate plot of average 2X differential pS187-H1.4 peaks near the promoter region before (Orange) and after (Blue) estradiol treatment. Following estradiol treatment, distinct enrichment of pS187-H1.4 over the untreated is observed. The signal was aligned +2Kb and −2Kb relative to the Refseq TSSs. CDK9 inhibitor treatment – Flavopiridol quenches the pS187-H1.4 signal (Green)

In order to further understand this enrichment pattern as a result of hormone induction, we next focused our study on the pS187-H1.4 peaks with at least a 2-fold increase following E2 induction.

### 2X differential pS187H1.4 signals enriched at promoters and can be quenched by CDK9 inhibition

In comparisons between ChIP in E2-treated and untreated MCF7 cells, we found 727 pS187-H1.4 differential peaks with at least a 2-fold increase in normalized mapped reads. We visualized this increase using an aggregate plot averaging the pS187-H1.4 signal across all differential peaks, to reveal a strong preferential enrichment at the promoter (Figure 3B - blue trace). The two-fold increase in pS187-H1.4 enrichment as a result of E2 treatment over the untreated control is pronounced, whereas the pan-H1.4 signal was depleted at the promoters, similar to the pattern described above.

Next, we tested the connection between CDK9 and H1.4 phosphorylation in this context, by ChIP-seq conducted after a 1-hour treatment with a CDK9-specific inhibitor, Flavopiridol (FLVP). After this treatment, the pS187-H1.4 promoter signal was significantly quenched (Figure. 3B - green trace). These data are consistent with a previous report by our lab, in which pS187-H1.4 was identified as a *bonafide* substrate for the kinase activity of CDK9 [9], and further verifies the pS187-H1.4 signal.

While the ‘metagenome’ profile and aggregate plots detect patterns of enrichment across the genome, they average the architecture of protein binding across all genes, and thus may obscure critical differences at the level of individual or subsets of individual loci. We used the GREAT platform (http://great.stanford.edu) to reveal the correlation of the nearest genes 1Kb upstream and downstream of the E2-induced pS187-H1.4 peaks for previously established functional annotations. The top ten most enriched annotations from the MSigDB perturbation category are listed with their enrichment p-values in Table 2. This ontology category contains gene sets that represent gene expression signatures of genetic and chemical perturbations [15]. As highlighted in the table, the strongest association of pS187-H1.4 peak-associated genes is with gene sets that were up-regulated as a result of estradiol addition in MCF7 cells, other breast cancer cell lines and even some breast tumor samples under a variety of conditions and with varying longer times of E2 treatments. This confirms that pS187-H1.4 strongly associates with promoters of bonafide E2 up-regulated genes.

**Table 2:**
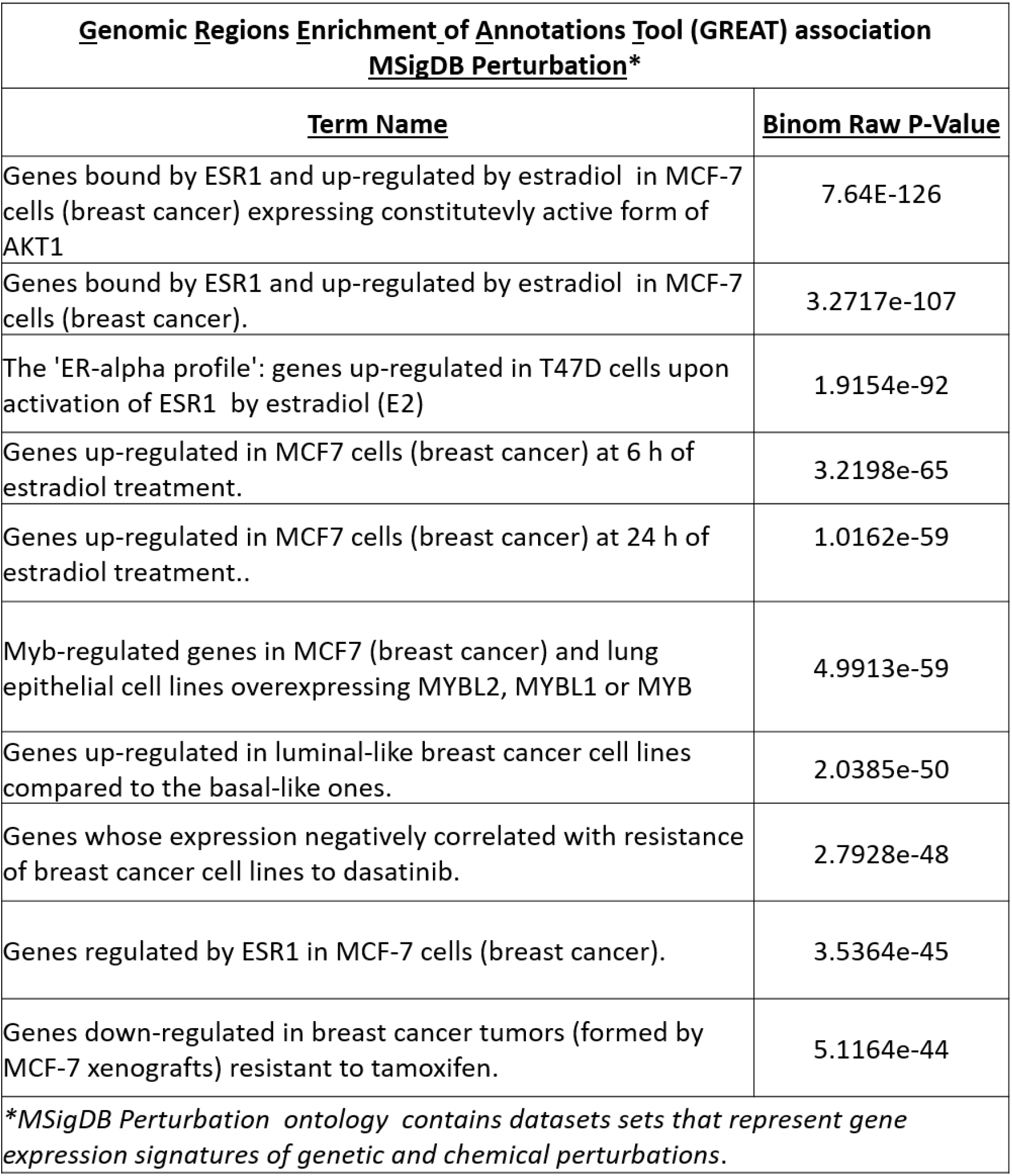
GREAT associations made with 2X differential pS187-H1.4 signal associated genes. Top ten categories in decreasing order of p-value listed.

| **<u>Genomic Regions Enrichment of Annotations Tool (GREAT) association</u>**<br>**<u>MSigDB Perturbation\*</u>** |  |
| --- | --- |
| **<u>Term Name</u>** | **<u>Binom Raw P-Value</u>** |
| Genes bound by ESR1 and up-regulated by estradiol in MCF-7 cells (breast cancer) expressing constitutively active form of AKT1 | 7.64E-126 |
| Genes bound by ESR1 and up-regulated by estradiol in MCF-7 cells (breast cancer). | 3.2717e-107 |
| The 'ER-alpha profile': genes up-regulated in T47D cells upon activation of ESR1 by estradiol (E2) | 1.9154e-92 |
| Genes up-regulated in MCF7 cells (breast cancer) at 6 h of estradiol treatment. | 3.2198e-65 |
| Genes up-regulated in MCF7 cells (breast cancer) at 24 h of estradiol treatment.. | 1.0162e-59 |
| Myb-regulated genes in MCF7 (breast cancer) and lung epithelial cell lines overexpressing MYBL2, MYBL1 or MYB | 4.9913e-59 |
| Genes up-regulated in luminal-like breast cancer cell lines compared to the basal-like ones. | 2.0385e-50 |
| Genes whose expression negatively correlated with resistance of breast cancer cell lines to dasatinib. | 2.7928e-48 |
| Genes regulated by ESR1 in MCF-7 cells (breast cancer). | 3.5364e-45 |
| Genes down-regulated in breast cancer tumors (formed by MCF-7 xenografts) resistant to tamoxifen. | 5.1164e-44 |
| *\*MSigDB Perturbation ontology contains datasets sets that represent gene expression signatures of genetic and chemical perturbations.* |  |

To visualize this estradiol induced pS187H1.4 enrichment at an individual gene, we used the UCSC genome browser [34] (Figure. 4A). We selected the Flotilin 1 (*FLOT1*) gene from the list of genes associated with the 2X pS187-H.14 differential peaks. Here we see, the *FLOT1* promoter is clustered very near the promoter of neighboring gene, *IER3*. Both promoters show a basal level of pS187-H1.4 signal in untreated cells; after E2 stimulation pS187-H1.4 ChIP signals were markedly increased. In addition to this increase in pS187-H1.4 signal after estradiol treatment, we also observed a clear overlap of the pS187-H1.4 signal with that of H3K4me3, as was seen genome-wide (Figure 1B). This pS187-H1.4 signal was quenched as a result of pre-treatment FLVP, as expected. Browser views of an additional representative gene, *TOB1* relative to housekeeping gene *ACTG1*, also demonstrated the increase in pS187-H1.4 signal as a result of estradiol treatment, the quenched pS187-H1.4 signal as a result of FLVP treatment and co-localization of active signal with RNAPII (Supplementary Figure. 7).

**Figure 4:**
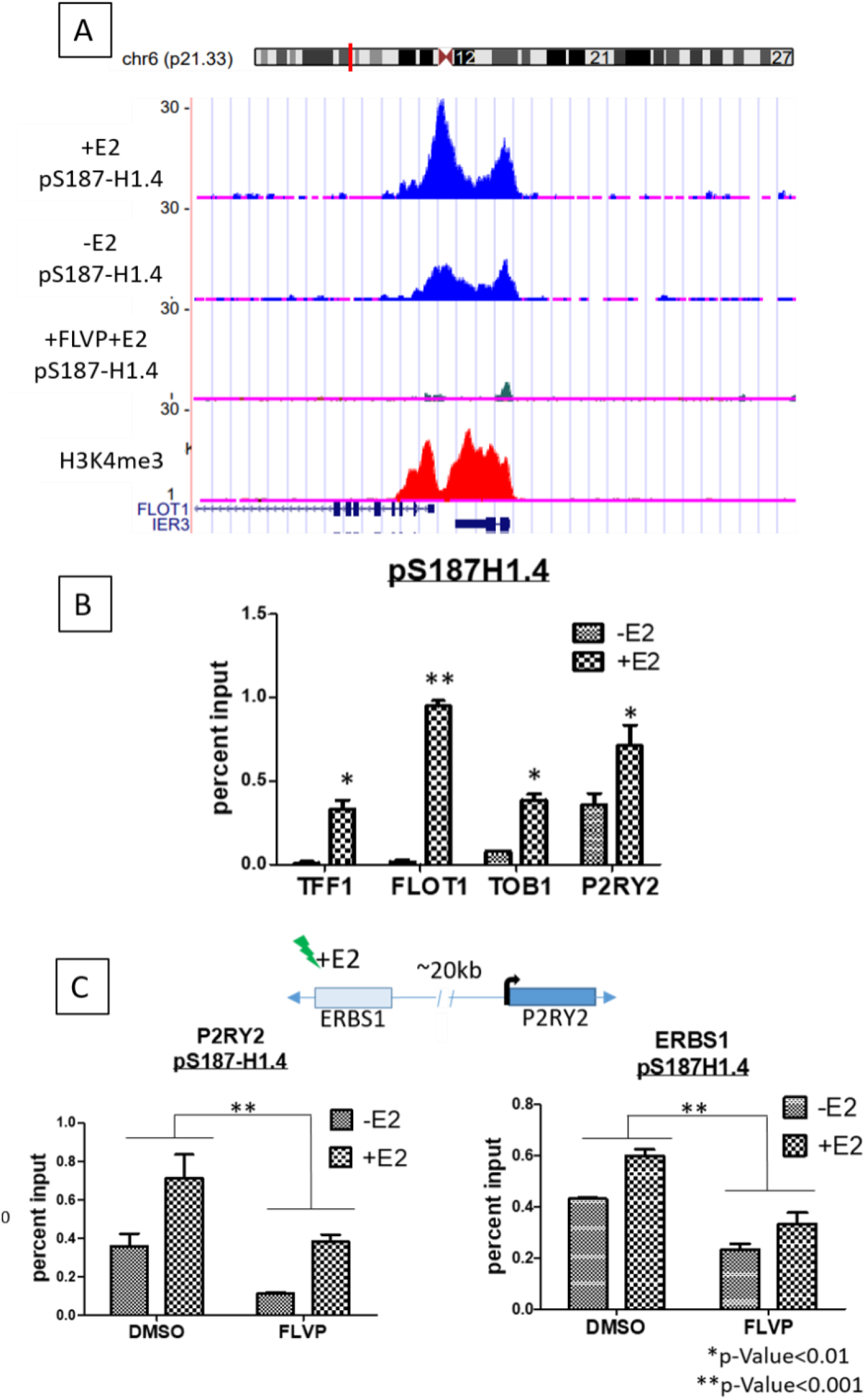
pS187-H1.4 2X differential signal verified on an individual gene level. (A) UCSC genome browser shot showing pS187-H1.4 signal before and after (−/+) estradiol treatment at FLOT1 and IER3 genes. The blue traces show pS187-H1.4 signals before and after estradiol treatment. The green track shows the pS187-H1.4 signal quenched as a result of FLVP treatment. H3K4me3 track in red is included to show ‘active’ state of genes. (B) ChIP-qPCR conducted with pS187-H1.4 antibody to verify changes on an individual gene level. Changes in pS187-H1.4 enrichment as a result of E2 addition shown at FLOT1, TOB1, TFF1 and P2RY2 promoters. All four genes show a significant increase in pS187-H1.4 enrichment as a result of E2 addition. (*p-value<0.01, **p-value<0.001) (C) ChIP-qPCR conducted with pS187-H1.4 antibody to show pS187-H1.4 enrichment at the P2RY2 gene promoter and its estrogen receptor binding site (ERBS1) that is ~20kb upstream of the promoter. The consequent loss of enrichment as a result of FLVP treatment is also shown. Rabbit immunoglobulin (rIg) used as a control for non-specific binding. Vehicle treated (DMSO) samples show an increase in pS187-H1.4 signal as a result of E2 addition. FLVP treated samples show an impaired ability of pS187-H1.4 enrichment. (*p-value<0.01, **p-value<0.001)

To confirm the genome-wide data, we performed ChIP-qPCR to verify and quantitate the E2-induced pS187-H1.4 increase on an individual gene level focusing on *FLOT1, TOB1*, *TFF1* and *P2RY2* (Figure 4B). We included *P2RY2* and its known estrogen receptor binding site, located approximately 20kb upstream of the promoter and ERBS1; based on previously published GRO-sequencing data these genes undergo transcription upregulation in response to E2 treatment [13, 17]. Primers overlapping the promoter regions of these genes were designed to test ChIP enrichment by quantitative PCR (qPCR). The promoters of *FLOT1, TOB1, TFF1* and *P2RY2* all showed a significant increase in the pS187-H1.4 ChIP signal at their promoters after E2 treatment (Figure 4B). To confirm and quantitate the loss of pS187-H1.4 signal as a result of FLVP treatment, *P2RY2* and *ERBS1* were used as a representative pair. The quenching of the signal observed as a result of CDK9 inhibition supports the idea that the signal observed was in fact that of pS187-H1.4 (Figure. 4C). In contrast, ChIP signal at housekeeping genes *ACTB* and *ACTG* remained relatively unchanged as a response to E2 treatment (Supplemental Figure. 5) although it was reduced as a result of FLVP treatment. Minor changes observed might be attributed to the involvement of these genes in cytoskeletal rearrangements as a result of E2 induction as previously described [35].

We also performed real-time qPCR (RT-qPCR) on candidate gene transcripts derived from this study to verify their individual gene expression levels after E2 exposure in our experimental system (Figure. 5). We confirmed that *TFF1, TOB1, TFF1* and *SMAD7* expression was significantly induced by E2, with a two-fold increase in signal. We further tested the direct role of H1 in the E2-activation of these genes by knocking down expression of H1.4 with RNAi. The induction of each of these E2-responsive genes was lost after H1.4 knockdown, suggesting that H1.4 is important for the E2 induced activation of these genes.

**Figure 5:**
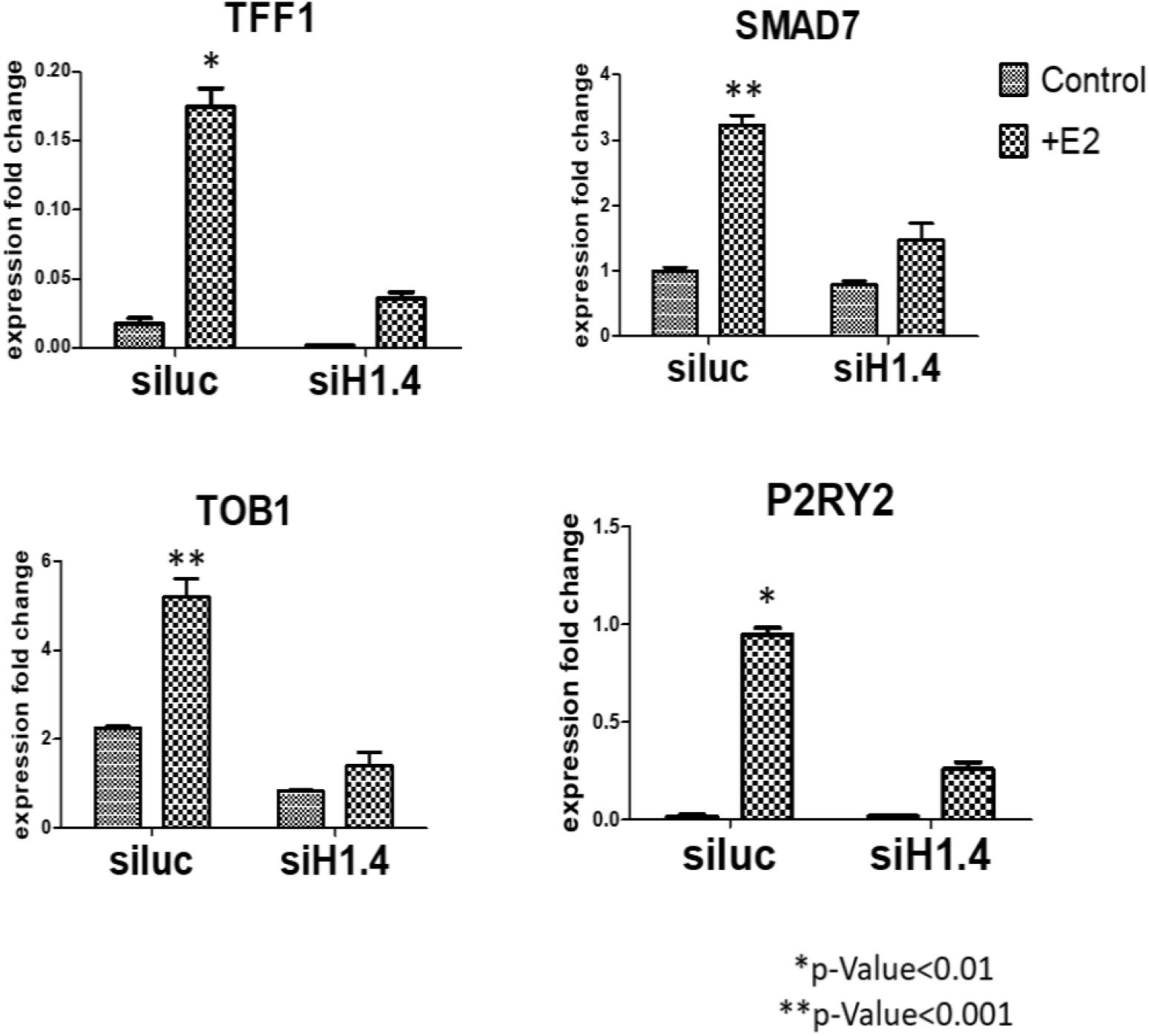
RT-qPCR to show importance of H1.4 in gene expression. Signals shown as a relative fold change in expression. Candidate genes from our ChIP-seq study were selected to show that E2 treatment led to an increase in gene expression. This coincides with the increase in pS187-H1.4 ChIP-PCR signal described in Figure.4B. *si*Luciferase (Control) was compared to *si*H1.4 to reflect loss of gene expression at H1.4 depleted genes. (*p-value<0.01, **p-value<0.001)

Taken together, the data presented here shows for the first time that a phosphorylated form of a linker histone variant i.e pS187-H1.4 is enriched at ‘active’ genomic regions, specifically promoters, when transcription is induced. This is in contrast to previous findings [2, 36] that reported a depletion of H1 by phosphorylation at transcriptionally ‘active’ sites.

### pS187-H1.4 enrichment is associated with previously identified estradiol responsive genes

In a study by Stendner et. al. 2007, 196 genes were identified as regulated by estrogen in MCF7 cells; the responsive genes were either stimulated or repressed by E2. This study also showed that transcription factor E2F was an early target for estrogen action and was a critical component of the hormone-induced proliferative response. To examine pS187-H1.4 enrichment at promoters of these early E2-responsive genes, we generated an aggregate plot of pS187-H1.4 signal around the promoters of these 196 genes (Figure 6). The pattern of pS187-H1.4 enrichment reproduced the trend that was observed in the genome-wide analysis of peaks as observed in Figure 3A, with the pS187-H1.4 enrichment at the promoter with the peak shifted towards the +1 nucleosome. The association of pS187H1.4 binding with verified early response elements - namely E2F binding sites - suggests that not only is phosphorylation of H1.4 associated with active promoters but may be involved in the early stages of estrogen-responsive gene activation.

**Figure 6:**
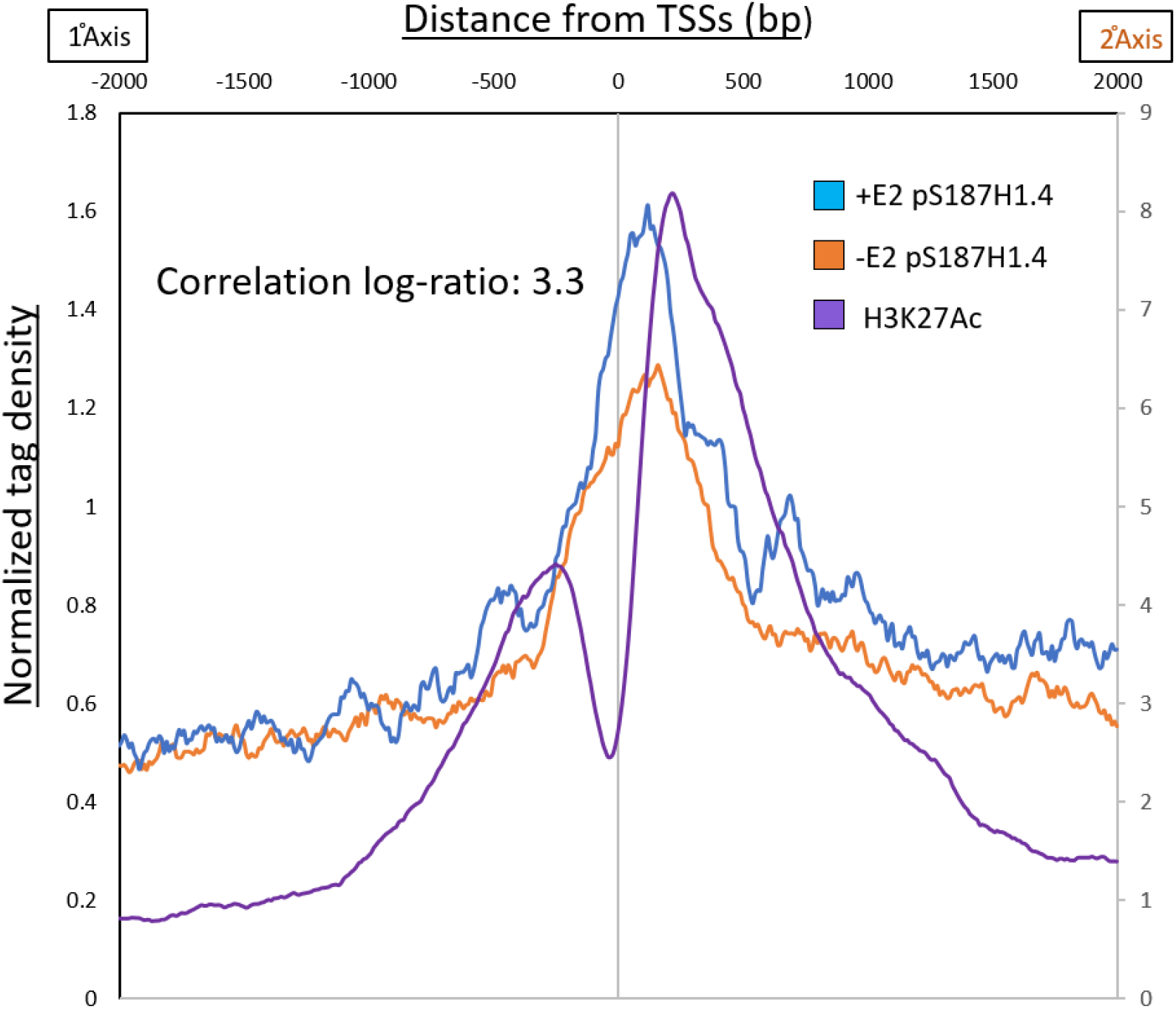
pS187-H1.4 at early responding E2 induced genes. 196 genes previously identified as being early targets of estrogen induction were analyzed for pS187-H1.4 signal at their promoters. Aggregate plot with average pS187-H1.4 signal before (orange) and after (blue) estradiol treatment plotted centered at refseq TSSs. A correlation ratio of pS187-H1.4 signal at these gene promoters was calculated as a log ratio (3.3)

## DISCUSSION

### pS187H1.4 enrichment at promoters marks active genes

Previous attempts to understand the functions of histone H1 have involved mapping the distribution of all H1 variants together across the genome, under the assumption that all of the variants were interchangeable and redundant in function. However, like core histones, H1 variants also undergo distinct post-translational modifications that impart different functionality to these proteins. Among the known modifications, phosphorylation is most abundant [2, 37], but the lack of specific antibodies has restricted the exploration of functions for these common H1 histone modifications.

In the present study, we have addressed this gap in knowledge by employing a unique antibody designed to detect a specific phosphorylated version of variant H1.4, pS187-H1.4. Early studies show that the sequence of human H1.4 shares a 93.5 sequence identity with its mouse ortholog [38, 39]; this level of conservation is much higher than was observed for other variants, indicating the functional significance of H1.4. Another report also pointed to the functional significance of this variant, demonstrating that the loss of H1.4 in T47D breast cancer cells resulted in cell death [40]. H1.4 undergoes various post-translational modifications (PTMs), such as K26 methylation [41] which allows binding of HP1 and subsequent heterochromatin function, and Gcn5-mediated H1.4K34Ac [42], which works to activate transcription by facilitating the binding of chromatin remodelers and recruitment of transcription factors. Furthermore, ChIP and qRT-PCR studies from our laboratory in Hela, NT2 and mESC cells have implicated pS187-H1.4 in transcriptional activation, showing enrichment of this modified variant at promoters of active genes [8, 9].

Although there have been a number of studies looking into histone PTMs, there is currently no genome-wide study describing the predominant interphase phosphorylation sites. This study provides the first genome-wide evidence of functions of pS187-H1.4, in the context of estradiol (E2) regulation. Earlier reports have described the estrogen response as being carried out through estrogen receptor isoforms (ERα or ERβ) dimerizing with 17β-estradiol and binding to specific motifs in the DNA to elicit a transcriptional response [43–45]. In this well-studied system of rapid signaling it is novel to observe that a variant of the linker histone - a family of protein usually associated with repression-is enriched at the transcription start sites of ERα/β-E2-activated genes. Data presented here strongly support this association and raise the intriguing possibility that the H1.4, in its phosphorylated form, may add another layer of transcriptional control that has not so far been identified. For example, surprisingly, pS187H1.4 binding is enriched in the nucleosome-depleted regions that are known to harbor active binding sites for transcriptional control intermediaries, co-regulators, and even RNAPII [13, 46, 47]. Especially given the observation of a significant, direct overlap of pS187-H1.4 and RNAPII across the genome, the possibility of direct/indirect association with other transcriptional regulators is now open for investigation.

In a previous study by Liao and colleagues [9] CDK9 was identified as the primary kinase responsible for the phosphorylation of S187H1.4. In this study, we have demonstrated that on a genome-wide level, the pS187-H1.4 signal at TSS is quenched upon treatment with the CDK9 inhibitor, Flavopiridol (Figure. 3). The CDK9 kinase is a part of the PTEF-b complex, which is responsible for the activation of RNAPII, by phosphorylating its CTD at Ser5, negative elongation factor (NELF) leading to its depletion. This is followed by Ser5 and DRB-sensitivity Inducing Factor (DSIF) phosphorylation, such that it becomes a positive elongation factor and travels with the polymerase [48]. Despite the obvious importance of DSIF and NELF as rate-limiting factors for PolII, growing evidence suggests that there may be additional factors may contribute to this regulation, including Gdown1 and TFIIF [49, 50]. With this study, we provide evidence that not only is pS187-H1.4 a substrate of the same kinase that is involved in the regulation of PolII, but it also significantly occupies the nucleosome-depleted regions as the ‘paused’ PolII throughout the genome. Together with evidence that pS187-H1.4 physically interacts with RNAPII and also appears to accumulate at a predicted +1 nucleosome location where ‘elongating polymerases’ are known to be found [51], we speculate that phosphorylated H1.4 may be a novel factor required in the ‘activation’ of transcriptional elongation or may even be involved in ‘ priming’ the chromatin to allow transcriptional elongation by RNAPII.

### Genome wide distribution of pS187-H1.4 reveals possible interaction with multiple transcriptional regulatory pathways

The association of pS187-H1.4 with features across the genome and particularly promoters, strengthens the hypothesis that H1.4 is important for the activation of genes via the RNAPII regulatory pathway. However, it is noteworthy to observe that there a strong enrichment of pS187-H1.4 observed in CpG Islands which are known to mainly contain sites of active transcription (Table 1). The regulation of these chromatin regions is noteworthy in that it depends on methylation state of the DNA and activity and /or recruitment of polycomb factors. In special relevance to this study, CpG islands are common sites of estrogen response elements (EREs) that are responsive to estradiol [52, 53]. We also observed that pS87H1.4 is strongly associated with non-coding RNAs (ncRNAs) and small nucleolar RNA (snRNA), both of which have different regulatory mechanisms in their response to gene activation stimuli. The role of ncRNAs in the form of lncRNAs like HOTAIR, MIAT, and H19 in estradiol response have only recently been described. In this context, the ncRNAs have been shown to regulate transcription by interacting with and guiding various chromatin modifying complexes like Polycomb-repressive complex 2 (PRC2) and Lysine-specific demethylase 1 (LSD1) [54]. Similarly, a distinct mechanism of activation is employed by snRNAs.

Together, this suggests that pS187H1.4 may be able to interact with multiple different transcriptional regulators, all with the final outcome of transcription activation. This suggests that globally pS187H1.4 may be a versatile factor with the capability of interacting with various transcription regulatory components. Although this study was focused on the estradiol-based response system as a model, we observed a basal level of pS187-H1.4 proximal to the promoters. Together these data suggest that pS187H1.4 may play a more global role in the regulation of transcription.

Since H1 has been largely regarded as a general repressor, the progress toward understanding the function and the mechanisms of action of individual H1 family members has been slow. Further, the paucity of appropriate ChIP-grade antibodies has also been a limiting factor. However, our novel, highly specific antibodies have allowed us to describe the role of pS187H1.4 in transcriptional activation for the first time, opening up a new area of study and suggesting that the roles of other specific H1 variants and their various modified forms should be further studied.

In order to further elucidate the effect of H1.4 phosphorylation on gene activation, the mechanism of action must be tested. In particular, the extent and nature of downstream effects must be assessed in cases where the phosphorylation event is prevented. Identifying the mechanism of CDK9 recruitment to this H1.4 will also yield an insight into the nature of its activity with regard to RNAPII regulation. However, these studies raise important issues regarding the functions of H1.4 variants that suggest broad significance in gene regulatory mechanisms, and which we are now well poised to address.

## AUTHOR CONTRIBUTIONS

CAM and AS conceived and designed the project. AS performed the experiments. CS performed the bioinformatics analysis. AS wrote the manuscript. LS, CS and AS edited the manuscript.

## DATA AVAILABILITY

ChIP-Seq data has been deposited in the GEO database with the accession number GSE137748.

## SUPPLEMENTARY DATA

Supplementary data available in a separate file.

## ACKNOWLEDGEMENT

We would sincerely like to thank Dr. Alvaro G. Hernandez and Chris L. Wright from the Roy J. Carver Biotechnology Center for performing and their guidance in the sequencing experiments and Huimin Zhang from Dr. Stubbs’ group for her help in library preparation. We would also like to thank Dr. Andrew Belmont, Dr. Benita Katzenellenbogen and Dr. Jongsook Kemper for their feedback on the project. We would also like to extend sincere thanks to Dr. Yamini Dalal for her support in the completion and submission of this manuscript.

