## Supplementary figures for "Site-specific phosphorylation of histone H1.4 is associated with transcription activation": Supplemental figures.pdf

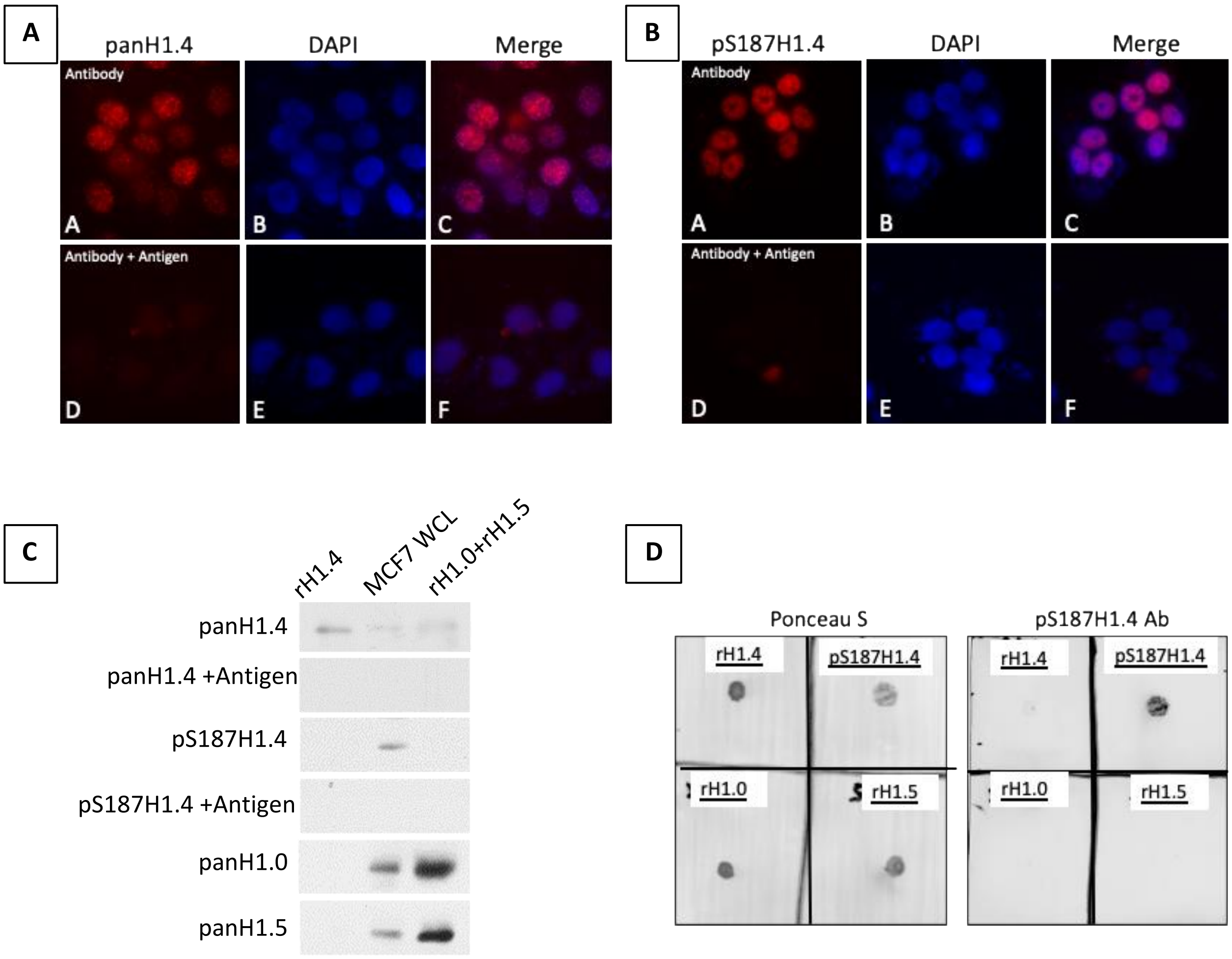


Supplementary figure 1. Antibody validation for pS187-H1.4 and pan-H1.4 affinity purified antisera. (A) Immunofluorescence performed on MCF7 cells to show pan-H1.4 staining (panels A-C). The pan-H1.4 signal was quenched when the immunofluorescence was performed with antigen-adsorbed pan H1.4 antibody (Panels D-F). (B) Immunofluorescent staining performed on MCF7 cells to show pS187-H1.4 staining (panels A-C). The pS187-H1.4 signal was quenched when the immunofluorescence was performed with antigen-adsorbed pS187-H1.4 antibody (Panels D-F). (C) Western blot performed with recombinant H1.4, MCF7 whole cell lysate (WCL) and a mixture of recombinant H1.0 and H1.5. pan-H1.4 and pS187-H1.4 antibodies used to show specific signal and demonstrate quenching when antibody was antigen adsorbed. (D) Dotblot performed to demonstrate specificity of pS187-H1.4 antibody. Left panel show ponceau staining of the recombinant H1.4, H1.0, H1.5 and pS187H1.4 peptide. The right panel shows specific staining with pS187-H1.4 antibody.

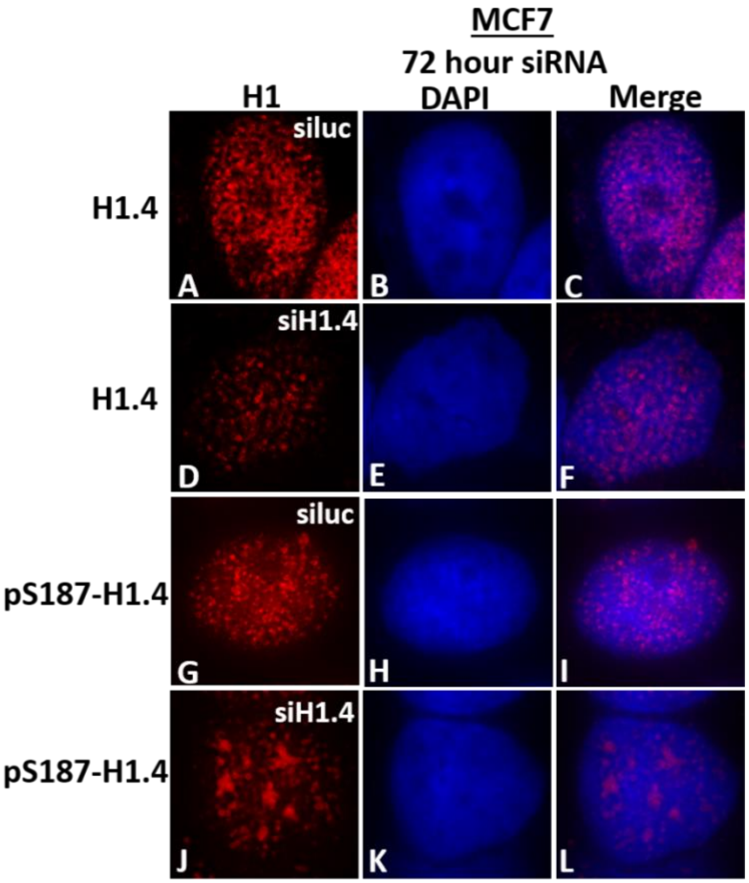


Supplementary Figure 2: Immunofluorescence of MCF7 cells (nuclei) treated with siLuc/siH1.4. Panels A-F shows pan-H1.4 staining of nuclei treated with siluc (A-C) and siH1.4 (D-F). Panels G-L shows pS187-H1.4 staining of nuclei treated with siluc (G-I) and siH1.4 (J-L).

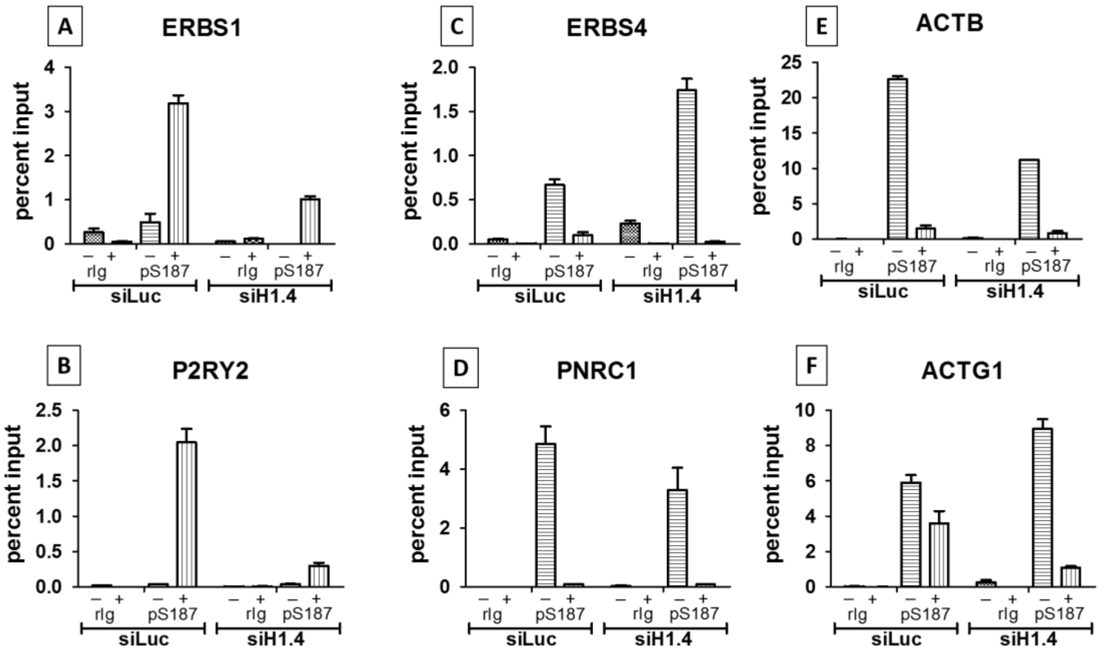


Supplementary Figure 3. Changes in the levels of pS187-H1.4 at promoters of genes responsive to estradiol (E2) (A-D) and housekeeping genes (E-F) as a result of siRNA treatment against H1.4 were assessed by ChIP-qPCR.

A-B) The levels of pS187-H1.4 at the promoters of P2RY2 and ERBS1 rise as a result of estradiol treatment. These levels drop as a result of H1.4 knockdown. (C-D) The levels of pS187-H1.4 at the promoters of PNRC1 and ERBS4 are repressed as a result of estradiol treatment. These levels diminish further as a result of siRNA mediated H1.4 knockdown. E-F) ACTB and ACTG1 appear to be repressed as a result of estradiol treatment. However, the pS187-H1.4 levels at the promoter of ACTB result in a further decrease of signal. pS187-H1.4 appears to increase as a result of the H1.4 knockdown but is reduced more than the siluc control when induced with estradiol. In all cases, negative control ChIP assays were employed non-immune rabbit immunoglobulin (rIg) in place of primary antisera.

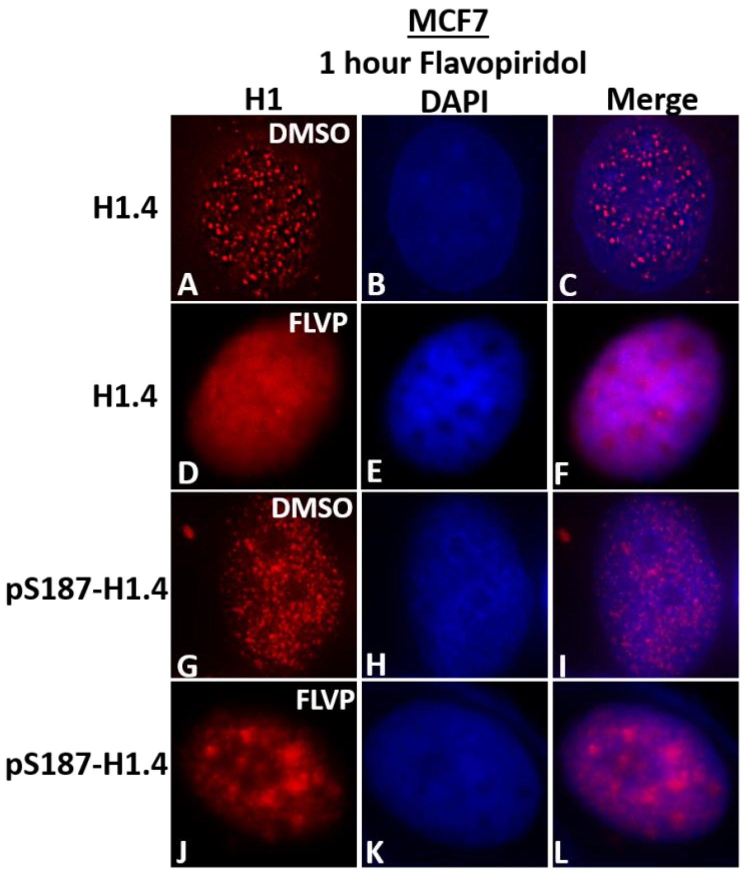


Supplementary Figure 4: Immunofluorescence of MCF7 cells (nuclei) treated with DMSO/FLVP for 1 Hour. Panels A-F shows pan-H1.4 staining of nuclei treated with DMSO (A-C) and FLVP (D-F). Panels G-L shows pS187-H1.4 staining of nuclei treated with DMSO (G-I) and FLVP (J-L).

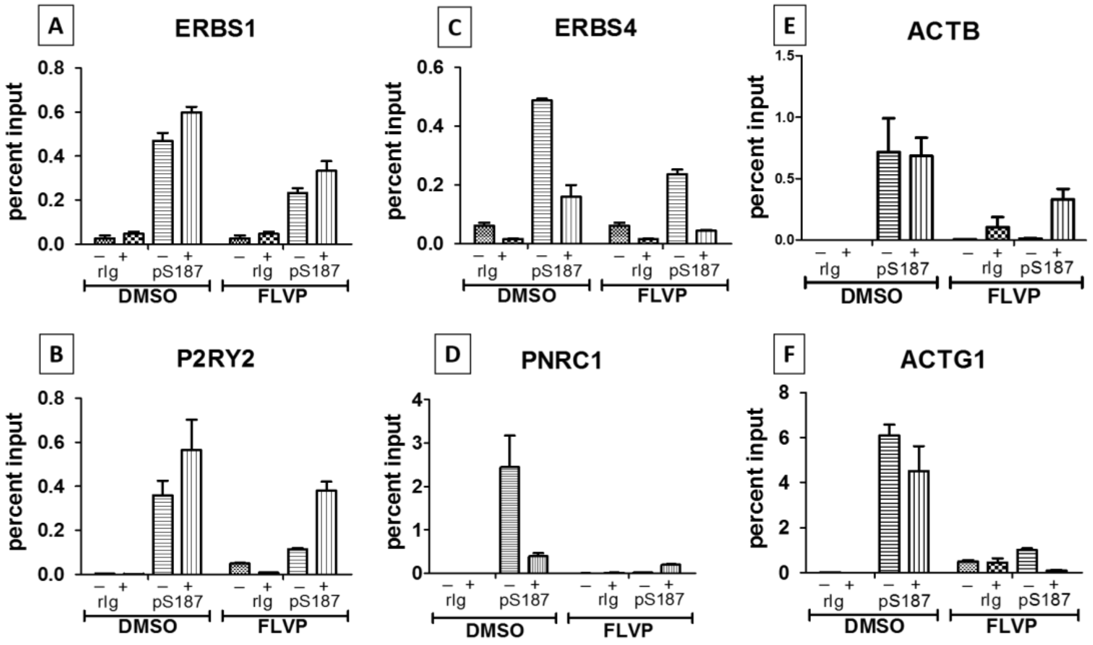


Supplementary Figure 5: Changes in the levels of pS187-H1.4 at promoters of genes responsive to estradiol (E2) (A-D) and housekeeping genes (E-F) as a result of FLVP treatment against H1.4 were assessed by ChIP-qPCR.

A-B) The levels of pS187-H1.4 at the promoters of P2RY2 and ERBS1 rise as a result of estradiol treatment. These levels drop as a result of H1.4 knockdown. C-D) The levels of pS187-H1.4 at the promoters of PNRC1 and ERBS4 are repressed as a result of estradiol treatment. These levels diminish further as a result of FLVP. E-F) ACTB and ACTG1 appear to be repressed as a result of estradiol treatment. However, the pS187-H1.4 levels at the promoter of ACTB result in a further decrease of signal. pS187-H1.4 appears to increase as a result of the H1.4 knockdown but is reduced more than the siluc control when induced with estradiol. In all cases, negative control ChIP assays were employed non-immune rabbit immunoglobulin (rIg) in place of primary antisera.

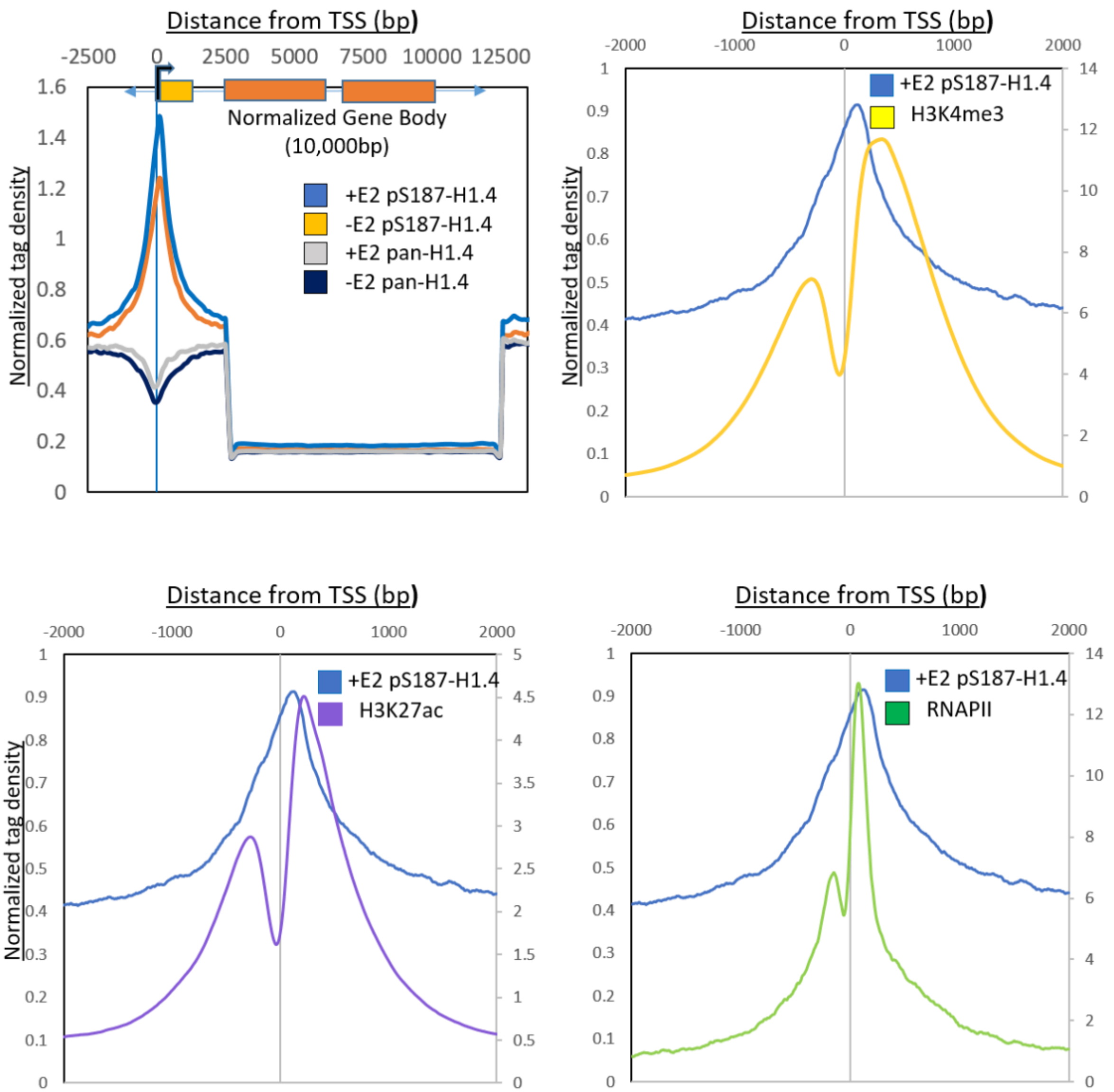


Supplementary Figure 6: Metagenome and aggregate plots demonstrating E2 induced pS187-H1.4 signal correlating with TSS and active transcription marks. (A) A metagenome profile generated with E2 induced pS187-H1.4 and pan-H1.4 ChIP-sequencing data to study enrichment across a typical gene. The gene body was mathematically defined and normalized to 10kb. +2.5kb and -2.5kb regions relative to the promoter were binned at 100bp. 2.5kb to 10kb represents a normalized gene body binned at 200bp. Typical transcription start sites (TSSs) showed maximum enrichment of the E2 induced pS187-H1.4 signal (blue trace). (B): Aggregate plot centered on the promoter showing average E2 induced pS187-H1.4 signals and its overlap with H3K4me3 signals. The signal was aligned +2Kb and -2Kb relative to the Refseq TSSs H3K4me3 was plotted on a secondary axis (label on right side). Overlap ratio: 3.25 (C): Aggregate plot centered on the promoter showing average pS187-H1.4 signals and its overlap with H3K27ac signals. H3K27ac was plotted on a secondary axis (label on right side). Overlap ratio: 4.99. (D) Aggregate plot centered on the promoter showing average pS187-H1.4 signals and its overlap with RNAPII signals. Overlap ratio: 3.14

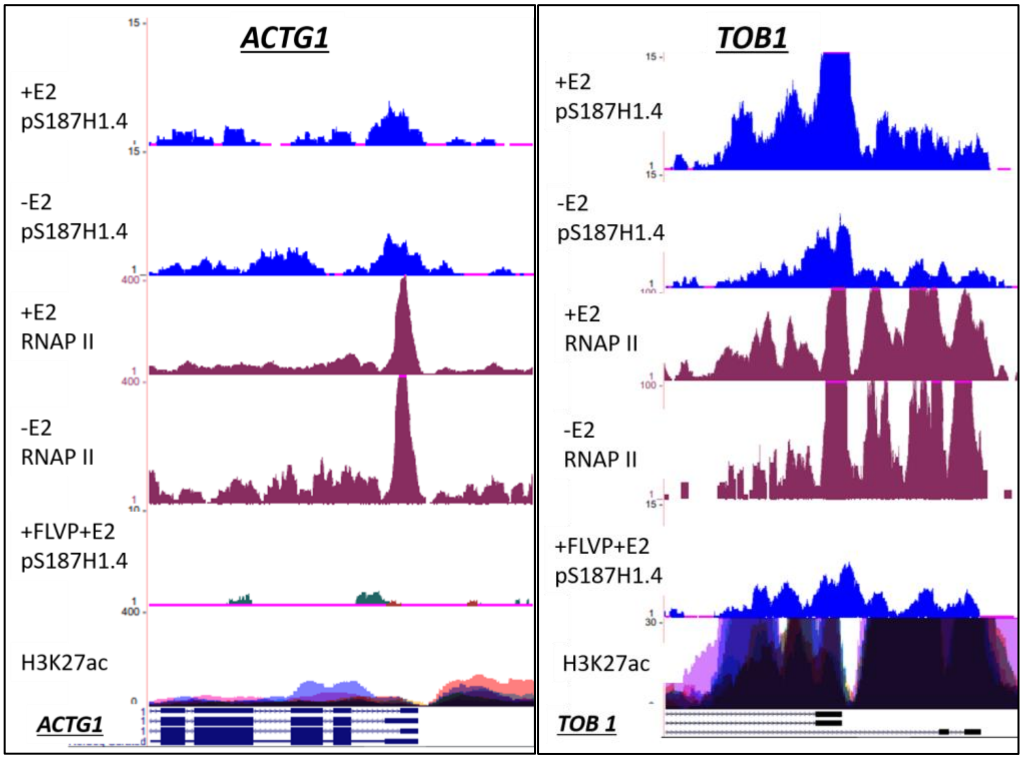


Supplementary Figure 7: UCSC genome browser shots of pS187-H1.4 signal (blue) before and after (-/+) estradiol treatment at mildly responsive housekeeping gene ACTG1 and fully responsive TOB1 genes and their co-localization with the RNAPII signals (magenta).

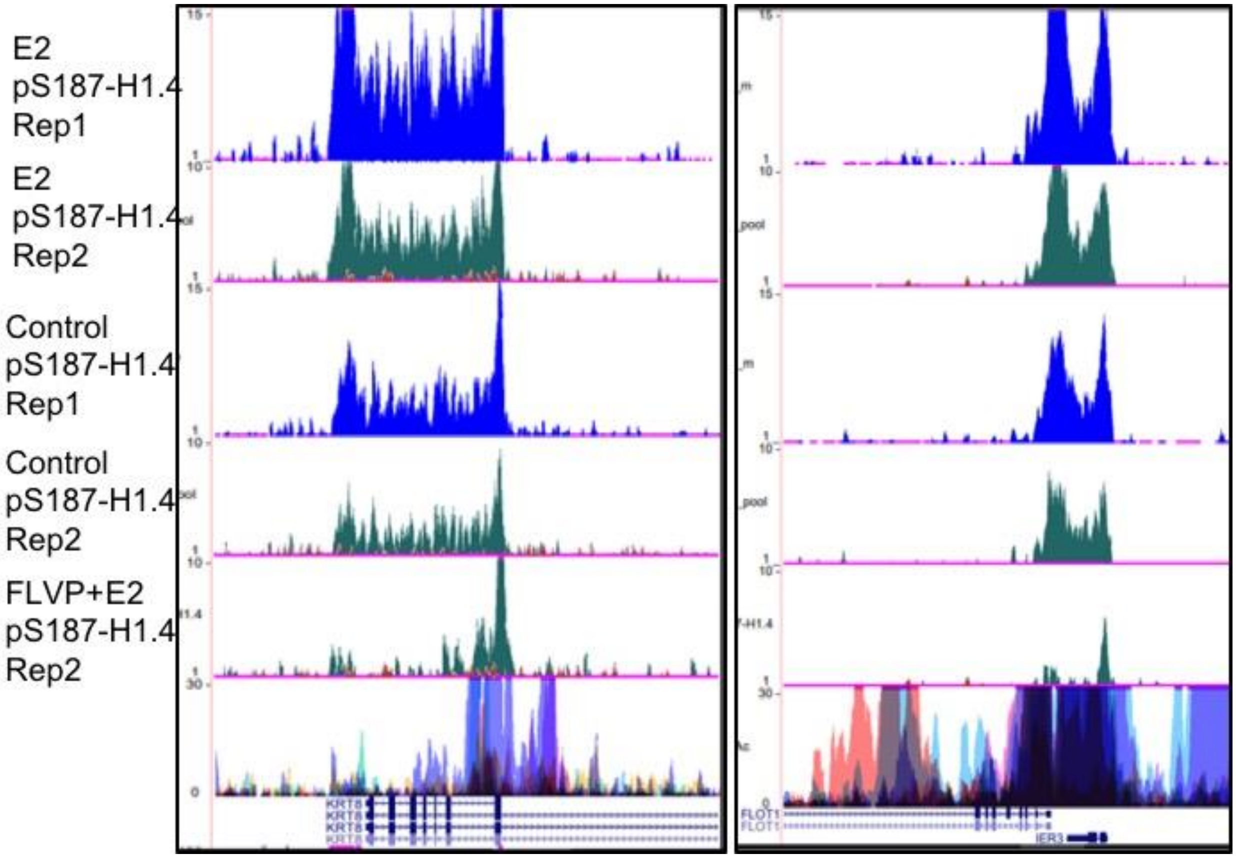


Supplementary figure 8:

Representative browser shots of two biological repeats of pS187-H1.4 signals before and after E2 treatment as well as pre-treatment with FLVP followed by E2, at individual genes. Repeat 1 is shown in blue and repeat 2 is shown in green. Trends observed are close to identical with an agreement value >0.94.

| Gene Name | Forward Primer | Reverse Primer |
|---|---|---|
| TFF1 | GAACAAGGTGATCTGCGCCC | CACTGTACACGTCTCTGTCTGG |
| FLOT1 | AAGCTGCCCCAGGTGGCAGAG G | TGTTCTCAAAGGCTTGTGATTCACC |
| TOB1 | GCTGTGTGGAGAAGTGAGCG | CTTGGGAGATCGCCGTTAGT |
| SMAD7 | TGGGTCCAAGGACAGATGTA | ACTCTCTGCATTGGTGAAGC |
| SOX2 | GGGGAAAGTAGTTTGCTGCC | GCTTAAGCCTGGGGCTC |
| POU5F1 | CTTCGCAAGCCCTCATTT | AGGTCCGAGGATCAACCC |
| ACTG1 | CGGCTTTCGGAAAGATCG | GAGCGGCGGAAGAACAGA |
| ACTB | GAAAGTTGCCTTTTATGGCTCG | TTACCTGGCGGCGGGTGT |
| P2RY2 | CGGTGGACTTAGCTCTGAGG | GCCTCCAGATGGGTCTATGA |
| PNRC1 | TCGCTCAGCAACGAAGAGAG | TCGCTCAGCAACGAAGAGAG |
| ERBS1 | AGGCAAATCCATTGTCATCC | AACTGGCTGGATCTTGAAGC |
| ERBS4 | GGCATAGCTAGGACCTCACC | GAGGGAGGAAAGTGGCTTCT |

Supplemental Table 1: Primers used for ChIP-qPCR
